# Proteome comparison between natural desiccation-tolerant plants and drought-protected *Caspicum annuum* plants by *Microbacterium* sp. 3J1

**DOI:** 10.1101/2020.03.23.004705

**Authors:** C García-Fontana, JI Vilchez, M Manzanera

## Abstract

Desiccation-tolerant plants are able to survive for extended periods of time in the absence of water. The molecular understanding of the mechanisms used by these plants to resist droughts can be of great value for the improvement of tolerance of sensitive plants with agricultural interest. This understanding is especially relevant in an environment that tends to increase the number and intensity of droughts. The combination of certain microorganisms with drought-sensitive plants can improve their tolerance to water scarcity. One of these bacteria is *Microbacterium* sp. 3J1, an actinobacteria able to protect pepper plants from drought. In this study we describe the proteome of the interaction of *Microbacterium* sp. 3J1 with pepper plants during droughts. We also compare this proteome with the proteome found in desiccation-tolerant plants. In addition, we studied the proteome of *Microbacterium* sp. 3J1 subjected to drought to analyze its contribution to the plant-microbe interaction. We describe those mechanisms shared by desiccation-tolerant plants and sensitive plants protected by microorganisms focusing on protection against oxidative stress, and production of compatible solutes, plant hormones, and other more specific proteins.

**IMPORTANCE:** Maintaining agricultural production under greater number and intensity of droughts is one of the main global challenges. Some plants can survive in the absence of water for extended periods of time. The molecular understanding of the mechanisms used by these plants to resist droughts is of great interest for the development of new strategies to face this challenge. Some microorganisms protect sensitive plants to some extent from droughts. *Microbacterium* sp. 3J1, is an actinobacteria able to protect pepper plants from drought. In this study we describe the different protein profile under drought used by the plant during the interaction with the microorganism and compare it with the one presented by desiccation-tolerant plants and with the one presented by *Microbacterium* sp. 3J1 to analyze its contribution to the plant-microbe interaction. We describe those mechanisms focusing on protection against oxidative stress, and production of compatible solutes, plant-hormones, and other more specific proteins.

## INTRODUCTION

Water scarcity is one of the most important limiting factors in agricultural production (1). Only a significant increase in food production can counterbalance the exponential growth of the population to supply enough food in the near future (2). To achieve this increase in food we have to ensure that crops are not lost due to more frequent droughts. Furthermore, we can recover previously abandoned farmland due to lack of water resources. We can use different approaches to grow plants for food purposes in these environments suffering from water limitation. One of these approaches consists of imitating the strategies followed by plants that naturally tolerate the lack of water, such as desiccation-tolerant plants and genetically modify the desiccation-sensitive plants (3). Another approach that have arised recently is the use of desiccation-tolerant microorganisms with the capacity to protect plants that would otherwise be sensitive to drought (4, 5).

Our group have identified a collection of desiccation-tolerant microorganisms, among them several actinobacteria stand out for their ability to tolerate desiccation due to the production of protective molecules called xeroprotectants (6–8). In addition, some of these microorganisms have the ability to colonize plant roots and protect these plants, such as pepper and tomato, against drought (9–11). A correlation between microorganisms with the largest capacity to protect plants from drought and their production of trehalose was found. *Microbacterium* sp. 3J1 was the microorganism with the highest production of trehalose and the one showing the highest protection of plants (11). By studying the metabolome of the interaction between the pepper plant root and *Microbacterium* sp. 3J1 under drought conditions, we have identified a change in the content of glutamine and α-ketoglutarate that resulted in the alteration of the C and N metabolism. As a result of this alteration, the concentration of sugars and amino acids was also modified due to the presence of the bacteria. In addition, antioxidant molecules, metabolites involved in the production of plant hormones such as ethylene, and substrates used for lignin production were altered in response to the presence of *Microbacterium* sp. 3J1 under drying conditions (12).

Similarly to desiccation-tolerant microorganisms there are some plants, such as *Xerophyta viscosa, Selaginella tamariscina, Craterostigma plantagineum, Boea hygrometrica*, able to withstand desiccation by arresting their metabolism during drying conditions and resuming such metabolism once water becomes available again (13). This process is termed anhydrobiosis and the plants are known as desiccation-tolerant plants or resurrection plants. To survive desiccation, these plants modulate the expression of a set of proteins involved in the decrease of photosynthesis, sugars accumulation, production of antioxidant molecules, or in the increase of the flexibility of cell walls and membranes (13, 14).

The study of proteomes is a versatile tool to understand the physiological processes involved in the tolerance of plants to absence of water. The study of proteomics instead of transcriptomics allows us to take into consideration transcript instability, post-transcriptional modifications and translational regulation of mRNA affecting gene expression. Changes in the quality and quantity of expressed proteins can be found with proteomic studies. To our knowledge, this is the first time the proteome of plants (*C. annuum*) protected by a microorganism (*Microbacterium* sp. 3J1) under drought conditions is described and compared with the proteome of naturally desiccation-tolerant plants. The comparison of both types of proteomes may form the basis for the development of alternative strategies to protect desiccation-sensitive food plants frequently affected by droughts.

## RESULTS AND DISCUSSION

### Experimental Set-up

We identified the interaction between *Microbacterium* sp. 3J1 and pepper roots for the protection of the plant against drought. Therefore, we decided to focus on the analysis of *C. annuum* in the presence and absence of the microorganism, in both cases under drought conditions. In addition, a comparison of the proteome profile of the 3J1 strain between drought and non-stressing conditions was performed to find out a potential contribution of *Microbacterium* sp. 3J1 to the pepper plant.

We decided to sample root material from 28-day old plants previously exposed to drought for 14-days to compare the proteome resulting from different microbiome compositions. *Microbacterium* sp. 3J1 was added to the plant 14 days after germination (Figure 1A). Then we subjected this material to the mapping of the roots proteome. In addition, the RWC of the plants was recorded, and total soluble protein was extracted and analyzed by 2D SDS–PAGE.

**FIGURE 1.**
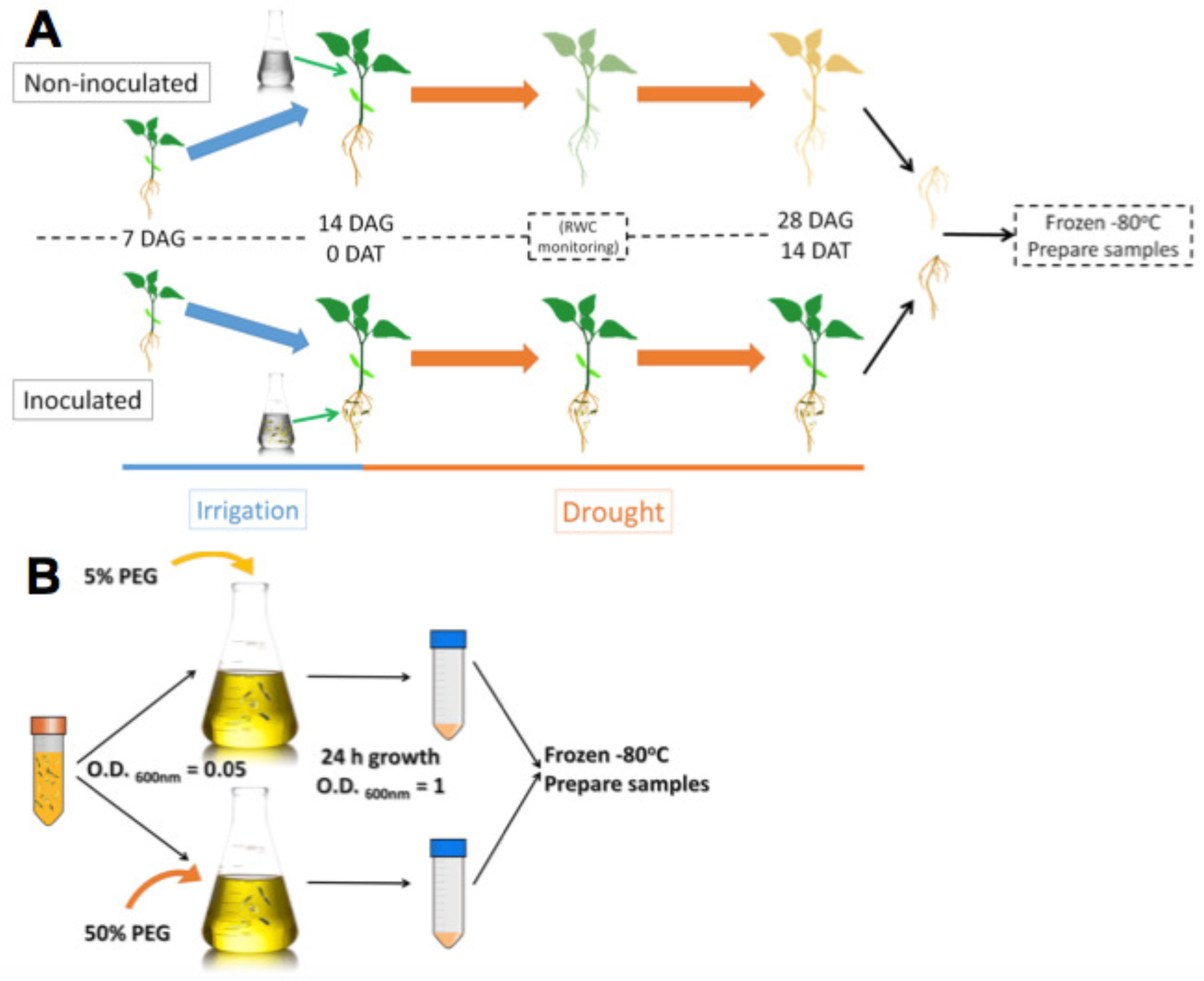
Root sample preparation workflow (A) and *Microbacterium sp.* 3J1 sample preparation workflow used in the proteomic study (B). DAG: days after germination; DAT: days after treatment. *Microbacterium* sp. 3J1-inoculated and non-inoculated pepper plant roots were sampled and preserved 14 DAT under drought conditions. Tests were performed with at least eight plants in triplicate. After freezing the samples, workflow continues with protein isolation and 2D-PAGE. *Microbacterium sp.* 3J1 cultures were grown until OD_600nm_ of approximately 1 in TSB supplemented with 5% or 50% PEG. Then samples were centrifuged and frozen until use. Protein isolation and 2D-PAGE were performed using the frozen samples.

For the *Microbacterium* sp. 3J1 proteome, cultures of the microorganism were supplemented with 5% and 50% polyethylene glycol (PEG), as indicated in the material and method section, to simulate the water activity of a non-stressed soil and of a soil subjected to drought-stress, respectively (Figure 1B).

### Analysis of differential proteins in *Microbacterium* sp. 3J1-inoculated and non-inoculated *C. annuum* roots under drought

After 14 days in the absence of watering, the RWC of non-inoculated plants was 0.4, while the RWC for inoculated plants was 0.68. These results show that plants were protected by the presence of *Microbacterium* sp. 3J1 (Figure 2). Inoculated and non-inoculated plants were analyzed by 2-DE to understand the proteome response of pepper plants subjected to drought to the presence of *Microbacterium* sp. 3J1. The root protein maps produced from three independent protein extractions showed a high reproducibility based on the analysis using the PDQuest software.

**FIGURE 2.**
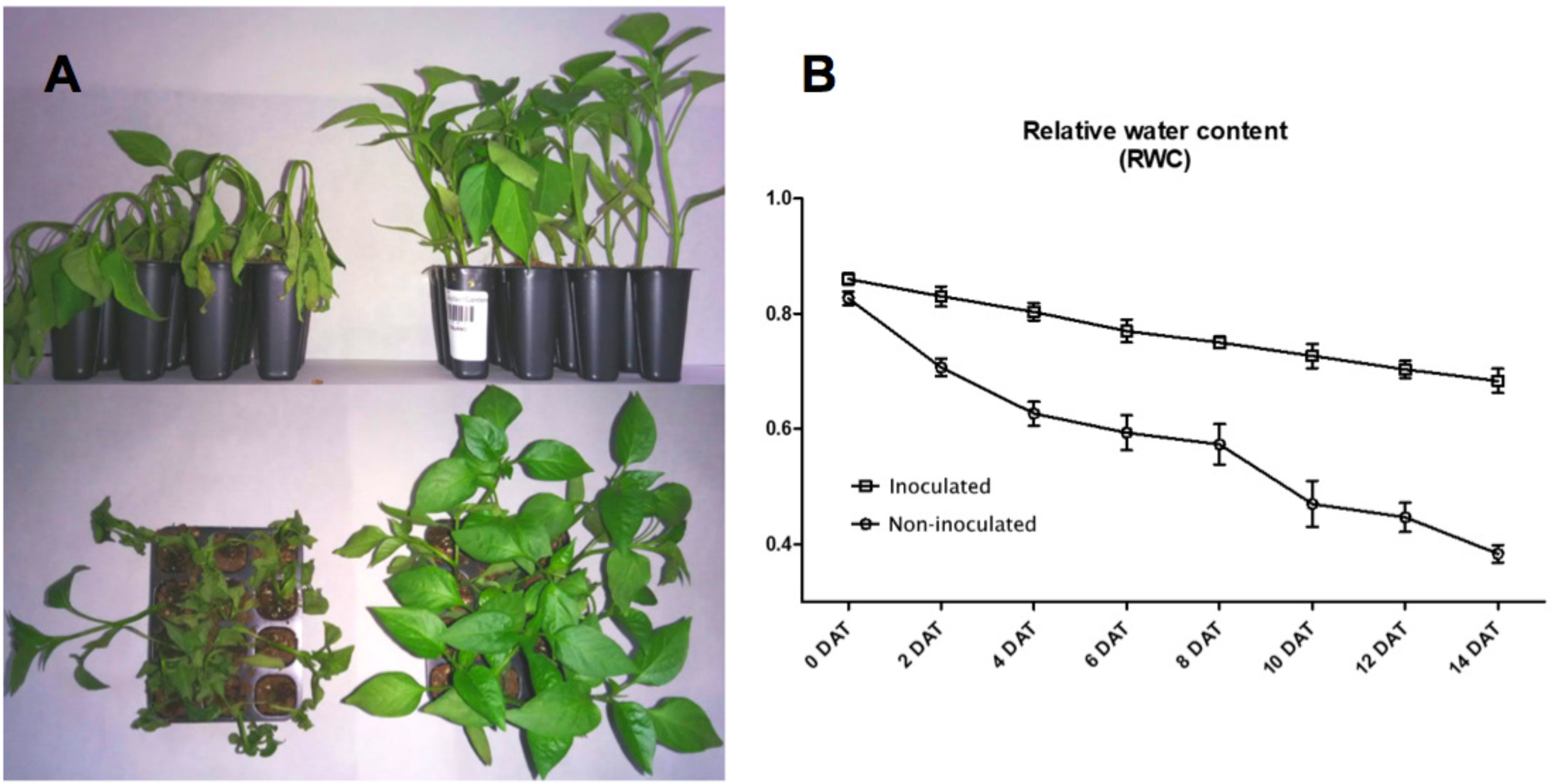
*Microbacterium* sp. 3J1-inoculated and non-inoculated *C. annuum* used in this study. **(A)** Phenotypes showed at 28 days after germination (DAG) of seedlings, at 14 days after treatment (DAT) with *Microbacterium* sp. 3J1 (left) and in absence of the microorganism as a non-inoculated condition (right), both in the absence of watering. **(B)** Drought critical point after treatment was determined by tracking the relative water content (RWC). Tests were performed with at least eight plants in triplicate. Population standard deviation (PSD) was used to determine the inner error bars.

Figure 3 shows representative gels of proteins extracted from the non-inoculated and inoculated plants. In conjunction, a total of 749 protein spots were reproducibly detected using PDQuest software from the non-inoculated samples and from the inoculated samples (*n* = 3). From a spot-to-spot comparison and based on statistical analysis, a total of 66 spots exhibited at least 2-fold (*p* < 0.05) difference in abundance between the non-inoculated and the inoculated plants (Figure 3A). Among 66 differential proteins, 30 spots showed qualitative changes (3 qualitative spots corresponded to inoculated-roots, whereas 27 spots were identified in non-inoculated roots. The rest 36 spots were present in both conditions, showing quantitative changes. A total of 27 spots out of these 36 differentially expressed spots were identified by MALDI TOF/TOF (Figure 3B, Table 1).

**TABLE 1.**
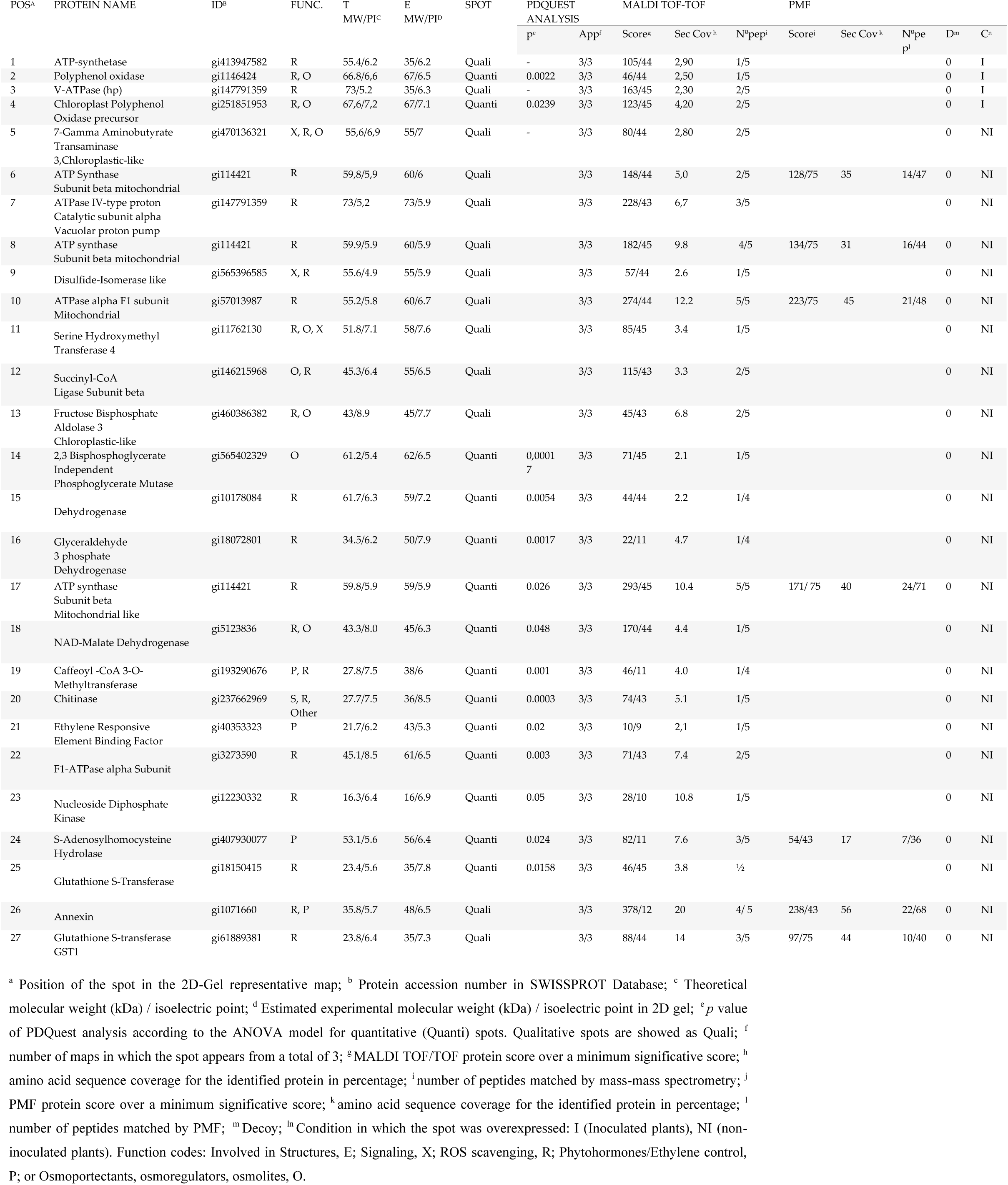
List of proteins from the proteome analysis showing significantly different amounts between inoculated and non-inoculated plants.

**FIGURE 3.**
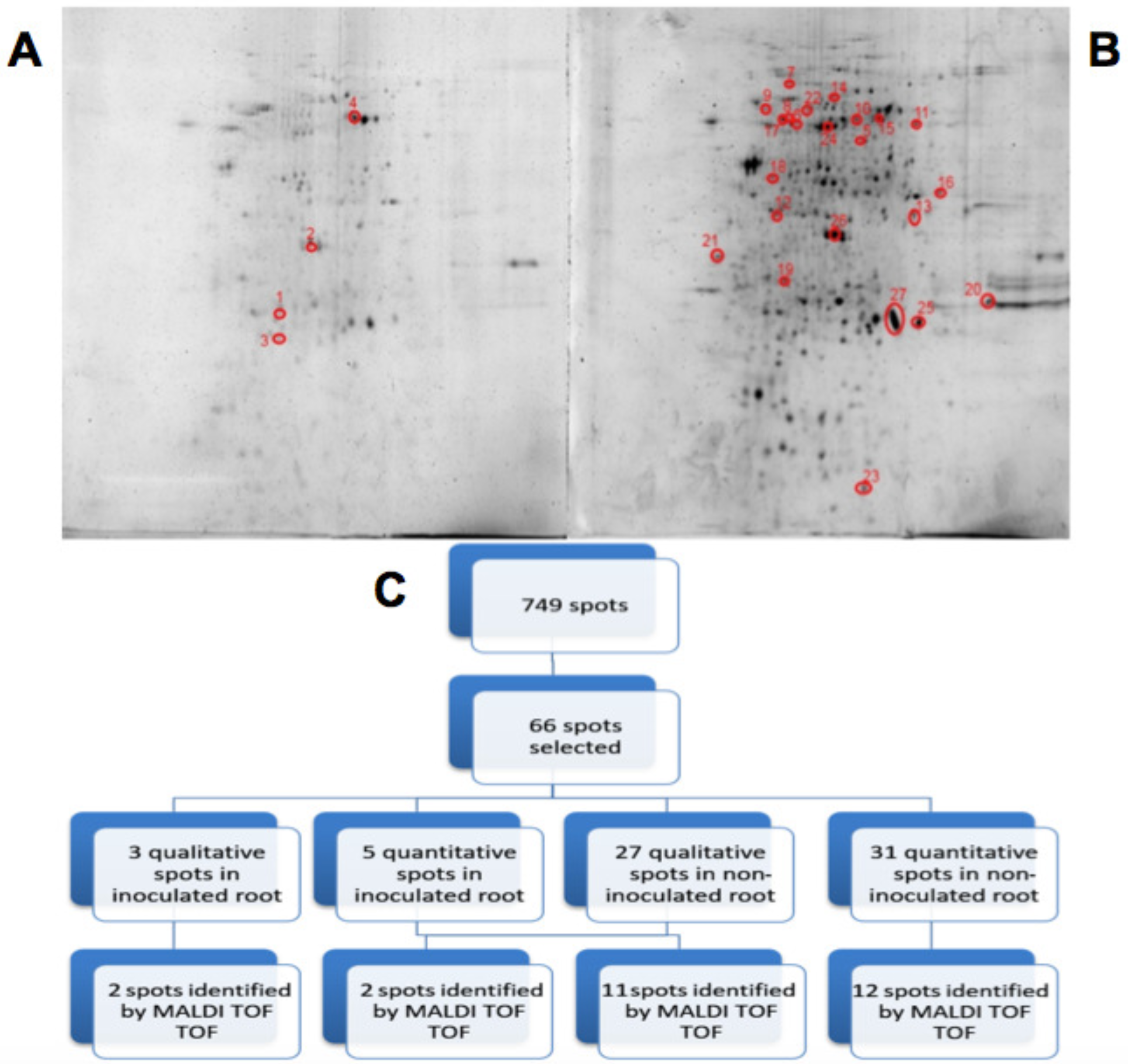
2D-PAGE image analysis of *Microbacterium* sp. 3J1-inoculated and non-inoculated plants subjected to drought. Upper pictures show differences in pepper root proteomes of non-inoculated (A) and inoculated with *Microbacterium* sp. 3J1 (B) seedlings obtained under drought conditions. Lower diagram shows the spot selection procedure and the final identified ones (C). Spots detection and selection were performed with PDQuest software v8.0. Red circles and numbers correspond to selected protein spots that were finally identified by MADI-TOF-TOF and described in Table 1. Pictures were selected as the most representatives from at least three replicates performed for each condition.

The number of protein spots detected for the *Microbacterium* sp. 3J1 protection of *C. annuum* against drought was within the range of protein spots described in previous studies on desiccation-tolerant plants. These plants included *Haberlea rhodopensis*, which showed a similar number of protein spots (152 *vs*. 148) when comparing hydrated control plants with those dehydrated (with 16% water content), showing 33 proteins specific for the desiccated *H. rhodopensis* plants only (15). For *Xerophyta viscosa*, 428 (±52) protein spots were identified using also the same software, although a lower difference in water content, changing from 65% to 35% RWC, was tested (16). In the case of *Boea hygrometrica*, the change in RWC from 100% to 2.4% was more drastic and a total of 223 protein spots were reproducibly detected (17).

### Proteins identified using MALDI-TOF/TOF-MS

To understand the function of differentially expressed proteins in *C. annuum* roots exposed to drought in the presence of *Microbacterium* sp. 3J1, the 66 differentially expressed spot proteins were analyzed by gel excision followed by a trypsin digestion and MALDI TOF/TOF-MS analysis in order to obtain the peptide mass fingerprint (PMF) data. Only 27 out of the 66 analyzed proteins could be identified (Table 1). These proteins were grouped in osmoprotectant producing proteins, reactive oxygen species (ROS) scavengers, plant-hormones, structural proteins, or signaling proteins. Similarly to the proteome of desiccation-tolerant plants exposed to dehydration, *C. annum* exposed to drought showed a diverse nature of differential proteins in the presence of *Microbacterium* sp. 3J1 compared to the non-inoculated plant. We consider this result to be indicative of the large impact of *Microbacterium* sp. 3J1 on plant cell function during dehydration. This is consistent with the diverse nature of differential proteins observed for desiccation-tolerant plants during dehydration processes.

### Osmoprotectans

As a result of the proteome analysis, we have identified the decrease in fructose bisphosphate aldolase and NAD malate dehydrogenase in *C. annuum* plants exposed to drought in the presence of *Microbacterium* sp. 3J1 (spots 13 and 18, respectively) compared to non-inoculated plants. Theses enzymes are involved in the production of glyceraldehyde 3-phosphate and oxaloacetate, respectively. Fructose-1,6-bisphosphate aldolase is a key metabolic enzyme that reversibly catalyzes the aldol cleavage of fructose-1,6-bisphosphate into dihydroxyacetone phosphate and glyceraldehyde-3-phosphate, either during the glycolytic pathway or gluconeogenesis and during the Calvin cycle (18). These reactions are involved in carbon fixation and sucrose metabolism and they are present in the chloroplast stroma and in the cytosol of green plants, allowing the plant to fix carbon dioxide into glyceraldehyde-3-phosphate, which can then be incorporated into other sugars (19, 20). The fact that the pepper plant produces increased amounts of these two enzymes points to the requirement of sucrose in the absence of *Microbacterium* sp. 3J1, whereas such requirement is no longer needed in the presence of this bacterial strain. We observed the accumulation of fructose (more than 8-fold), fructose-6-phosphate (approximately 18-fold increase), and trehalose (approximately 85-fold increase) in a recent metabolomic characterization of the pepper plant subjected to drought in the presence of *Microbacterium* sp. 3J1, compared to the non-inoculated plant. This result points to an alternative use of sugars as more efficient osmoprotectant (12). The concentration of other sugars was also increased in response to the microorganism when pepper plants were subjected to drought. Similarly, one of the mechanisms that desiccation-tolerant plants use to respond to drought consists of the accumulation of osmoprotectants, osmoregulators and osmolyte substances. For instance *X. viscosa, C. plantagineum, Sporobolus stapfianus*, and other resurrection plants modulate their carbohydrate metabolism for the production of sucrose (16, 21–23). However, *Selaginella lepidophylla* accumulates oxaloacetate, fumarate, succinate, and alpha-ketoglutarate (intermediates of the tricarboxylic acid cycle) during dehydration, therefore the importance of these metabolites for the metabolic flux through these pathways during dehydration or in preparation for rehydration is suggested (23). Nevertheless, trehalose is a minor component in a few desiccation-tolerant angiosperm species despite its importance in desiccation tolerance among lower organisms (24). Despite the differences in the profile of osmoprotectants produced by desiccation-tolerant plants and by *Microbacterium* sp. 3J1-inoculated plants, Ingle et al. have described the increase in the abundance of proteins (phosphopyruvate hydratase) involved in the control of glycolysis-gluconeogenesis pathways in the desiccation-tolerant plant *X. viscosa* (16). Oliver et al. also described the increase in enzymes associated with glycolysis including aldolases, phosphoglycerate kinase, and glyceraldehyde-3-phosphate dehydrogenase in *S. stapfianus* subjected to drying conditions (25). All the increased proteins: phosphopyruvate hydratase (in *X. vicosa*), phosphoglycerate kinase and glyceraldehyde-3-phosphate dehydrogenase (in *S. stapfianus*), and fructose biphosphate aldolase and NAD malate dehydrogenase (in *C. annuum* plants exposed to drought in absence of *Microbacterium* sp. 3J1) would allow an increase in flux through the gluconeogenic route during drought, providing an increased pool of hexose phosphate substrates to deal with the higher demand of sucrose in both types of plants (natural desiccation-tolerant plants and 3J1-protected pepper plants) under drying conditions.

### Energy metabolism

The addition of *Microbacterium* sp. 3J1 to pepper plants subjected to drought resulted in the production of two proteins (spots 1, 3) not found in non-inoculated plants, the disappearance of three proteins (spots 6, 7 and 10) and the decreased in the amount of two proteins (spots 17, 22) related to the production of ATP. This change in the expression pattern of proteins involved in the production of ATP may produce a different cell concentration of ATP. The availability of ATP during drought is critical because this metabolite is needed for the production of osmoprotectants. The increased amount of certain proteins involved in the production of ATP substituting other ATP producing proteins has been reported for several resurrection plants such as *B. hygrometrica* (17), *S. stapfianus* (25), *X. viscosa* (16), *Tortula ruralis* (26), and for *S. tamariscina* (27), where the increased production of ATPases was suggested to play a role in protein folding or unfolding and protein degradation in response to drought stress. Despite previous studies have reported that phosphorylation of high mobility group proteins reduced their binding to DNA, inhibiting replication and transcription in drought-sensitive plants such as maize (28), we have described the potential role of DNA as xeroprotectant, i.e. as dehydration protectant (29). Therefore we do not rule out a potential role of the over production of ATP in DNA production for DNA repair and, we do not discard that ATP can function as xeroprotectant by the production of DNA in the plant. The increase in proteins involved in DNA related processes have been reported for some desiccation-tolerant plants, such as *S. stapfianus* (25). This increase suggest a shared mechanism between desiccation-tolerant plants and *C. anuum* protected by *Microbacterium* sp. 3J1.

### Oxidative metabolism

The proteome analysis of *Microbacterium* sp. 3J1-inoculated *C. annuum* plants subjected to drought is marked by the decrease in abundance of proteins associated with oxidative metabolism, and more specifically, with enzymes involved in the protection against reactive oxygen species (ROS), pointing to a protection against ROS by alternative methods. ROS causes deleterious effects on the respiratory system, metabolic pathways, genomic stability, membranes and organelles during drought events. The increased production of ATP occurs in conjunction with the protection against ROS, and in this sense, we have found increased production of certain ATP producing enzymes as above described. Glutathione protects against oxidative damage by quenching activated oxygen species and participates in the metabolism of hydrogen peroxide (30). In addition to those proteins, we have identified an increased production of proteins involved in glutathione production such as glutathione-S-transferase and glutathione S-transferase GST1 (spots 25, 27). In the absence of *Microbacterim* sp. 3J1, the plant produces higher quantities of ROS scavengers, showing a special need to fight ROS, something that does not seem to be required in the presence of the 3J1 strain. The production of ROS scavengers is a common feature among desiccation-tolerant plants subjected to drying conditions (25, 31–34). Jiang and coworkers identified increased production of glutathione S-transferase in response to dehydration in *B. hygrometrica*, in special during the early stages of dehydration using proteome assays (17). The participation of glutathione to protect desiccation-tolerant plants has been described in many species, including *H. rhodopensis* (35), *B. hygrometrica* (17), *S. stapfianus* (25), to name a few. Apart from glutathione-producing enzymes, other proteins such as ascorbate peroxidase (ascorbate acts as a scavenger of hydrogen peroxide), or superoxide dismutase have been reported as proteins involved in oxidative mechanisms in several desiccation-tolerant plants, such as *X. viscosa* (36, 37). However, we have only found a decrease in the proteins needed for the production of glutathione for this type of antioxidant proteins in response to the presence of *Microbacterim* sp. 3J1.

### Cell signaling

An appropriate signaling system and the production of regulatory molecules, such as plant hormones, are needed for a coordinated response between the microorganism and the plant for its protection against drought. Therefore, we would expect the detection of proteins involved in the production of such molecules. A decrease in four proteins involved in the production of plant hormones was found in *Microbacterium* sp. 3J1-inoculated plants. These four proteins are caffeoyl-CoA 3-O-methyltransferase, ethylene responsive element binding factor, S-Adenosyl homocysteine hydrolase and annexin (spots 19, 21, 24, 26 respectively). Caffeoyl-CoA 3-O-methyltransferase (spot 19), catalyzes the conversion of S-adenosyl-L-methionine (SAM) and caffeoyl-CoA into S-adenosyl-L-homocysteine (SAH) and feruloyl-CoA, thus reducing the concentration of SAM. The production of ethylene in plants is mainly based in the presence of SAM, and therefore a decrease in caffeoyl-CoA 3-O-methyltransferase would translate into a reduction of ethylene and a concomitant release from senescence of the plant due to lack of water. The S-adenosyl homocysteine hydrolase (spot 24) is therefore involved in the reversible production of S-adenosyl homocysteine, contributing to the reduction of SAM and, consequently, reducing the ethylene production. The production of both enzymes has been described to increase in the drought-tolerant plant *C. dactylon* (38). The SAM to SAH ratio is involved in the enhancement of drought tolerance where methylation is a key step (39). In addition, we have previously described a reduction in the production of ethylene in pepper plants subjected to drought when inoculated with *Microbacterium* sp. 3J1 (12). This is of special relevance since we have observed the decrease in the ethylene responsive element-binding factor (spot 21) in pepper plants subjected to drought when *Microbacterium* sp. 3J1 was present. In resurrection plants, such as *X. viscosa*, it has been suggested that ethylene could be a signal for their tolerance to drought through the expression of *XVT8* (a gene encoding a glycine-rich protein, with significant identity to dehydrins) (40). However, the abscisic acid (ABA) is a more common plant hormone for the response to drought in *X. viscosa* (16). ABA controls the stomata closure and the accumulation of metabolites, increases the activity of antioxidant enzymes and the expression of genes encoding protective proteins in response to dehydration and it has been described as a major component in the hormonal control of the acquisition of desiccation tolerance in *S. stapfianus* (41), and in other resurrection plants (25, 42). The existence of a non-ABA signaling pathway has been proposed for *S. stapfianus*, where brassinosteroids and methyl jasmonate could play a role (43, 44)

### Polyphenol metabolism

We found an increase in the production of polyphenol oxidase (PPO) and the chloroplastid precursor of PPO (spots 2 and 4, respectively) in plants subjected to drought in the presence of *Microbacterium* sp. 3J1. An increase in PPO has been described to prevent of proteolytic activity for the conservation of the proteins during the desiccation state in *Ramonda servica* and *B. hygrometrica* (17, 45). The PPO has also been described as ROS scavenger in *H. rhodopensis, M. flabellifolia*, and *Ramonda serbica* (46–48). A potential role of PPO in Mehler-like reaction for detoxification of reactive oxygen species has been suggested because most PPOs are localized in the chloroplast thylakoid lumen; in addition, the chemical structure of PPOs makes them ideal for free radical scavenging activities in the chloroplast (49, 50). Therefore, we suggest that the presence of *Microbacterium* sp. 3J1 in pepper plants subjected to drought use PPO instead of glutathione to prevent the damage produced by ROS.

### *Microbacterium* sp. 3J1 proteome in response to drought

In order to assess the potential impact of the proteins produced by *Microbacterium* sp. 3J1 on the pepper plant during drought, we characterized the differential protein expression of this soil microorganism in response to severe drought by addition of PEG. The addition of PEG results in the sequestration of water molecules reducing the hydric potential of the media. To simulate the change from a non-stressed soil to a severe drought-stressed soil, liquid cultures of *Microbacterium* sp. 3J1 were supplemented with 5% and 50% (w/v) PEG 8000, respectively. The 2-DE was performed following the previously described conditions. A fold-change of at least 2 was considered as the minimum level of differentially expressed proteins when analyzing the 2-DE images. A total of 237 protein spots were analyzed as statistically significant (*p*-value <0.05) and 43 protein spots were selected and analyzed by MALDI-TOF/TOF-MS-(Figure 4). A total of 40 quantitative (19 up- and 21 down-regulated) proteins and 3 qualitative spots were differentially expressed proteins between 5 and 50% PEG conditions. Among these 43 analyzed spots, 13 were successfully identified (Table 2).

**TABLE 2.**
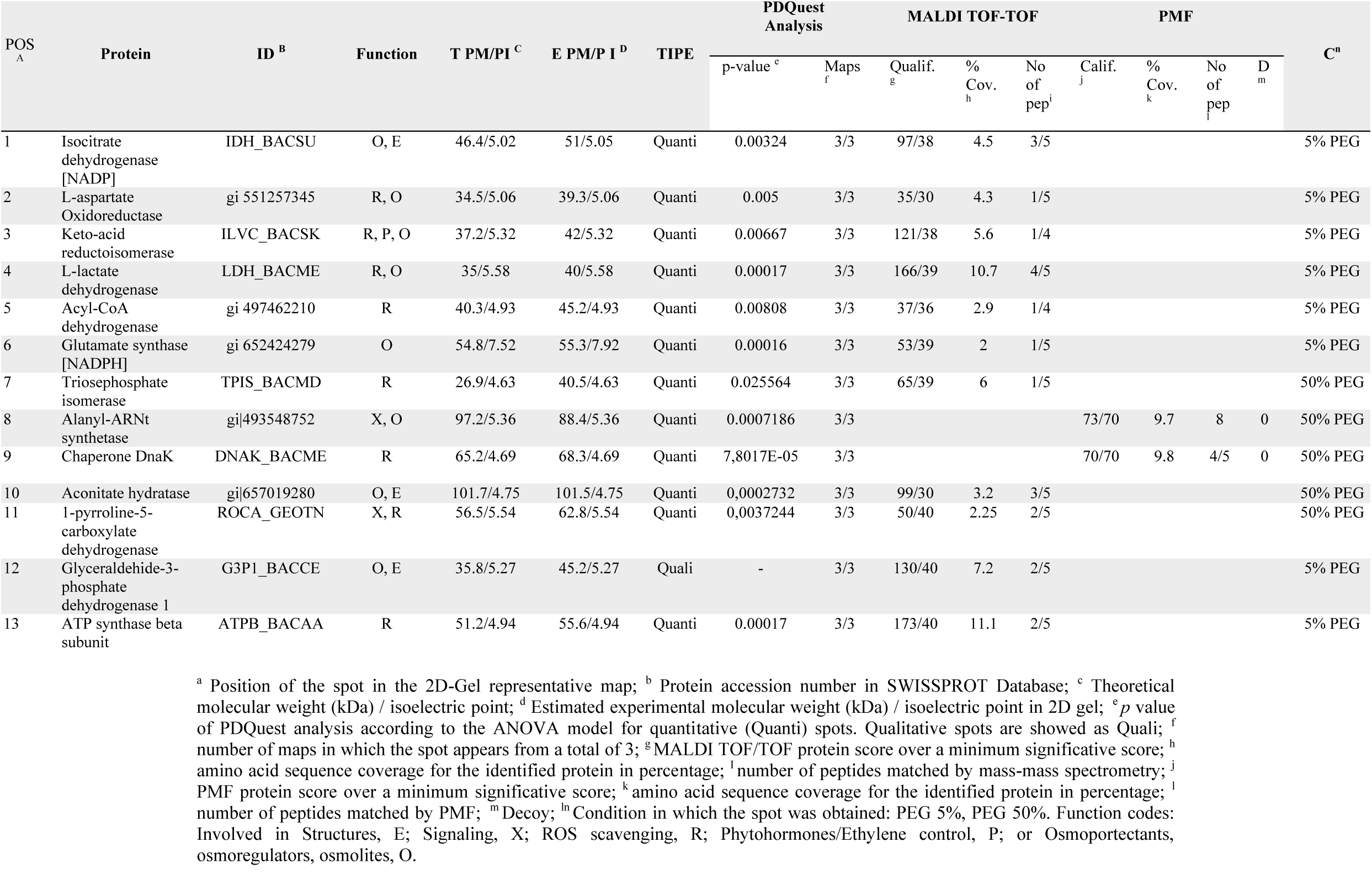
List of proteins from *Microbacterium sp.* 3J1 proteome analysis showing significantly different amount between 5% and 50% PEG-grown cultures.

**FIGURE 4.**
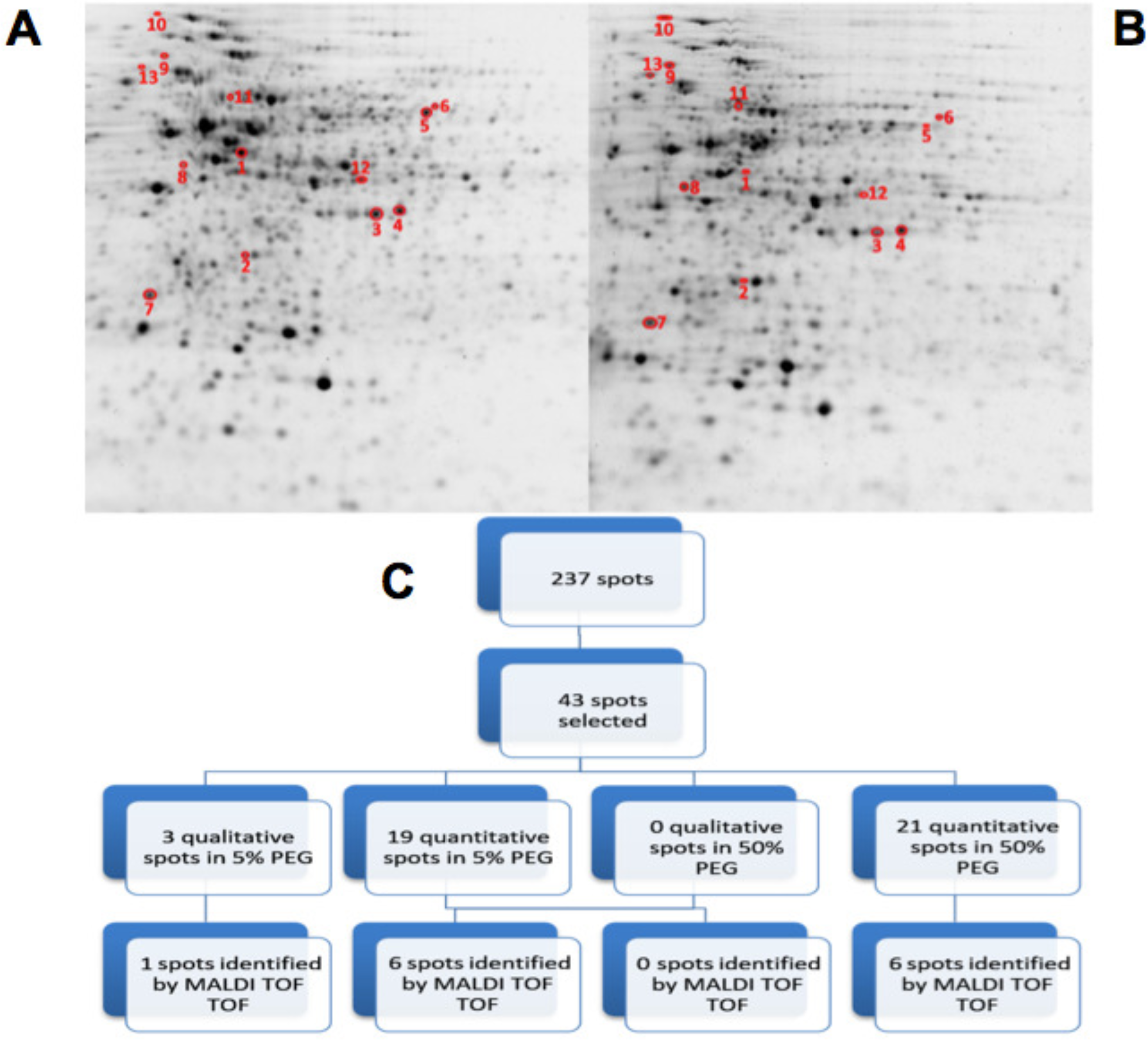
2D-PAGE image analysis of drought-subjected *Microbacterium sp.* 3J1 cultures. Upper pictures show differences in *Microbacterium sp.* 3J1 proteome when 5% or 50% PEG (B) were supplied to TSB growth medium. Lower diagram shows the spot selection procedure and the final identified ones (C). Spots detection and selection were performed with PDQuest software v8.0. Red circles and numbers correspond to selected protein spots that were finally identified by MADI-TOF-TOF and described in Table 2. Pictures were selected as the most representatives from at least three replicates performed for each condition.

All differentially identified proteins were analyzed via the NCBI BLAST database and the function of these proteins was explained by inputting the Uniprot accession number into the Uniprot database as shown in Table 2 (51, 52). The identified proteins were mainly involved in (a) osmotic protection, (b) energy metabolism, (c) antioxidant protection and (d) cell signaling.

Regarding the proteins involved in osmotic protection, 8 proteins were differentially expressed (spots 1-4, 6, 8,10, 12), including the decreased expression of L-aspartate oxidoreductase in media supplemented with 50% PEG (spot 2) compared to medium with 5% PEG. This enzyme is involved in the conversion of L-aspartate into oxaloacetate, resembling the catalytic step produced by the increased expression of malate dehydrogenase observed in *C. anuum* plants subjected to drought in the presence of *Microbacterium* sp. 3J1. In the presence of water stress conditions, we have also found a decreased production of NADPH-isocitrate dehydrogenase (spot 1) and an increased production of aconitase (spot 10), involved in the citric acid cycle, for the reversible catalytic conversion of citrate to isocitrate via *cis*-aconitate. The decreased production of NADPH-isocitrate dehydrogenase that catalyzes the irreversible conversion of isocitrate into α-ketoglutarate in conjunction with the increased production of aconitase in response to water stress conditions would be translated into a reduction of α-ketoglutarate by the partial inversion of the first steps of the citric acid cycle. The metabolomic analysis of the pepper plant subjected to drought also showed a dramatic decrease in α-ketoglutarate due to the presence of *Microbacterium* sp. 3J1. This decrease seems to respond to a decreased production of α-ketoglutarate and to the conversion of α-ketoglutarate into glutamine for the drastic change in the C and N metabolism of the plant due to the presence of the microorganism (12). In the inoculated plant subjected to drought, the concentration of glutamine is increased compared to the non-inoculated plant; however, no alteration in the glutamate was found. This might be explained by the observed decreased production of NADPH-glutamate synthase (spot 6) in *Microbacterium* sp. 3J1, when exposed to drying conditions (50% PEG). The produced glutamate from glutamine can be transformed into proline, a well described osmoprotectant, by enzymatic and non-enzymatic transformation. The last of the enzymatic steps requires the participation of the 1-pyrroline-5-carboxylate dehydrogenase, one of the increased proteins in *Microbacterium* sp. 3J1 in response to higher water stress (spot 11). The production of both proline and aspartate contribute to the production of osmoprotectants. Glutamate metabolism for the production of other amino acids such as alanine, valine and isoleucine, could be reduced also by the observed decreased production of keto-acid reductoisomerase (spot 3) under water stress. An increased production of alanyl-tRNA synthetase (spot 8) was found. This is a key enzyme for the availability of amino acids also overproduced by *Microbacterium* sp. 3J1 in response to water stress. A disappearance of glyceraldehyde-3-phosphate dehydrogenase (spot 12) was also found in response to water stress in *Microbacterium* sp. 3J1. This enzyme catalyzes the reversible conversion of glyceraldehyde-3-phosphate into 1,3-bisphosphoglycerate, contributing to the metabolic flux of the glycolysis-gluconeogenesis pathways described in the inoculated peppers in presence of *Microbacterium* sp. 3J1. This could also be the case of the increased production of triose phosphate isomerase (spot 7) found in *Microbacterium* sp. 3J1 under water stress, since this enzyme catalyzes the reversible conversion of glyceraldehyde-3-phosphate into dihydroxyacetone phosphate, allowing the availability of free sugars for the production of osmoprotectant, as described for 3J1-inoculated pepper plants under drying conditions.

A decreased production of the L-lactate dehydrogenase (spot 4) was observed in response to water stress. This protein reversibly catalyzes the inter-conversion of pyruvate and lactate, which may serve for the connection of the altered glycolysis-gluconeogenesis pathways with the altered citric acid cycle, therefore allowing an increased production of glucose.

Finally, an increased production of the chaperone protein DnaK (spot 9) was found in response to water stress by increasing the PEG concentration in the media from 5% to 50%. The DnaK protein actively participates in the response to hyperosmotic shock and it is also involved in chromosomal DNA replication. As previously described by our group, an increased production of DNA by *Microbacterium* sp. 3J1 was found in response to drought, where the DNA molecules seem to have a role as osmoprotectact as proved *in vitro* for the protection of proteins against desiccation (29).

An increased production of the β subunit of the ATP synthetase (spot 13) was found in response to water stress. We associate this increased production with the energy metabolism of *Microbacterium* sp. 3J1 in an analogous manner to the need of ATP during droughts in plants. Another finding was the decreased production of acyl-CoA dehydrogenase in response to water stress (spot 5). Acyl-CoA dehydrogenases are enzymes catalyzing the *α, β*-dehydrogenation of acyl-CoA esters in fatty acid and amino acid catabolism. We associate this reduced production of the acyl-CoA dehydrogenase under drying conditions with being part of the shut down of the lipid metabolism within the general arrest of metabolism found in anhydrobionts during the desiccated state. A reduced production of the acyl-CoA dehydrogenase was observed in *Bradyrhizobium japonicum*, a desiccation-tolerant microorganism, in response to other stressors, such as acidic pH (53), or in *Acinetobacter baumannii* and in the cyanobacteria *Leptolyngbya ohadii* among other desiccation-tolerant microorganisms in response to desiccation (54, 55).

## CONCLUSION

The desiccation-tolerant bacteria *Microbacterium* sp. 3J1 has been characterized at biochemical and metabolomic levels due to its response to drought. Despite the ability of *Microbacterium* sp. 3J1 to protect a diverse range of plants from drought, its response to drought was not characterized at the proteomics level to date. The ability to tolerate desiccation has been described in several microorganisms, and many of them have been described as able to protect plants from drought. However, the information available on the proteomics of desiccation-tolerant microorganisms and their role in protecting plants from drought is scarce. In the present study, we observe a parallel pattern between the protection of plants from drought conducted by *Microbacterium* sp. 3J1 and the mechanisms that naturally desiccation-tolerant plants use to survive desiccation. Despite of the large variety of mechanisms used by plants, a common strategy consisting of increasing the production of osmotic protectants, increasing energy metabolism, improving the antioxidant protection and reshaping the cell signaling in the plant via plant hormone production was found. We describe a similar pattern in pepper plants protected from drought by *Microbacterium* sp. 3J1, where all these molecular strategies are altered by an altered expression of proteins. The presence of *Microbacterium* sp. 3J1 in pepper plants subjected to drought seems to respond to the mechanism that the bacterium itself uses to survive drought, where the microorganism seems to take control of the plant metabolism for a higher chance of survival. We believe that this knowledge may form the basis for the development of alternative strategies to protect desiccation-sensitive food plants such as *C. annum*, which are affected by droughts frequently.

## MATERIALS AND METHODS

### Bacteria and Plant growth conditions

Green Pepper *Capsicum annuum* L. cv. Maor plants were produced by the specialist grower SaliPlant S.L. (Granada, Spain). Plants were grown in green houses under constant relative humidity (50–60%). The plants were illuminated with a 12-h light/dark cycle. Once plants were purchased, they were incubated in the lab under similar humidity and light/dark cycle. For the light cycle, 200 μmol photons·m^-2^·s^-1^ were used. In addition, dawn–dusk cycles were included with 150 μmol photons·m^-2^·s^-1^. The temperature ranged from 18 to 20°C during the dark period and from 20 to 25°C during the light period.

For inoculation, half of the plants were treated with 40 mL of bacterial suspension (10^8^-10^9^ CFU/mL) in sterile saline solution, while the other half of the plants were used as non-inoculated controls and watered with the same volume of sterile saline solution.

Plants were regularly watered and allowed to acclimate for at least 2 weeks after inoculation before simulation of drought was implemented.

For inoculation, *Microbacterium* sp. 3J1 was used as a rhizobacteria drought-tolerance enhancer. These bacteria were grown in tryptic soy broth (TSB) at 30°C and 180 rpm (56).

### Dehydration treatment and determination of the relative water content (RWC)

Three groups of *C. annuum* plants, each consisting of three individual plants, were dried down by withholding water. A control group of plants was watered with an equivalent volume of saline solution in the absence of inoculant. Plants of each condition (with and without inoculant) were extracted from the substrate and completely cleaned of soil by several washes in distilled water. Roots tissues were cut and preserved until sample preparation and protein extraction by wrapping in aluminum foil and storing at −80°C after treatment with liquid nitrogen.

Fresh weight (FW), dry weight (DW), and fully turgid weight (FTW) of the whole plants free from soil were recorded. The relative water content (RWC) was calculated according to Vílchez et al. (2016) and to Mayak et al., (2004) using the following equation RWC = (FW-DW) × (FTW-DW)^-1^. In addition, root length (RL) and stem length (SL) were measured.

Polyethylene glycol 8000 was added to the microorganism culture to simulate dehydration. *Microbacterium* 3J1 were grown in tryptic soy broth (TSB) supplemented with 5% (w/v) PEG 8000, at an initial absorbance of 0.05 at 600 nm wavelength. The cultures were incubated for 24 h and then split into two. Half of cultures were supplemented with 50% PEG 8000 to simulate severe drought. The cells were further incubated at 30°C and 180 rpm until stationary phase was reached and then, harvested by centrifugation at 15,000 g for 30 min. The cell pellets were stored at −80°C until use.

### Isolation of total protein

Frozen pellets, consisting of 0.5 g of plant tissue or cells collected from 10 ml of *Microbacterium* sp. 3J1 cultures were resuspended in 1 mL lysis buffer containing 500 mM Tris-HCl pH 6.8, 4% sodium docecyl sulfate (SDS) (w/v), 30% glicerol (v/v), 1 mM EDTA, 5 mM phenylmethanesulfonyl fluoride (PMSF), 200 mM dithiothreitol (DTT), 0,01% Benzonase® Nuclease (v/v) (Roche).

For the plants and bacterial pellets, cells were broken using FastPrep (three cycles of 40 s at 6.5 m/s). The lysate was centrifuged for 15 min at 20,000 g and 4°C, and the supernatant was recovered for soluble protein quantification by Bradford Method. Soluble proteins (150 μg) were precipitated. Protein precipitation was performed by adding 50% (v/v) ice cold trichloroacetic acid (TCA) to obtain a final concentration of 10% TCA, which was then incubated on ice for 15 min. The samples were centrifuged for 15 min at 20,000 g and 4°C, and then the protein pellet was washed three times with 1 ml of chilled acetone and it was centrifuged at 4°C for 5 min at 20,000 g. Finally, the dry samples were stored at −20°C until use.

### Two-dimensional polyacrylamide gel electrophoresis (2D-PGAGE)

Total soluble protein was dissolved in 300 μL of rehydration buffer (RH) (8 M Urea, 2 M Thiourea, 2% (w/v) Chaps, 0.5 mM PMSF, 20 mM DTT, 0.5% (v/v) Bio-Lyte Ampholyte (BioRad) and trace amounts of bromophenol blue, at 15°C 2h and 200 rpm. The protein samples were then centrifuged for 15 min at 20,000 g and 4°C.

Supernatants, containing 150 μg soluble protein fraction, were separated by isoelectric focusing (IEF) using a 17-cm nonlinear pH 3.0-10.0 immobilized pH gradient (IPG) for plant proteins and a nonlinear pH 4.0-7.0 IPG strips (ReadyStrips, BioRad) for bacterial proteins in the first dimension. IEF was performed at 20°C. Strips were first actively rehydrated at 50 V for 14 hours which accumulated to 288 V/h. IEF was performed using a Protean i12 IEF System (BioRad) with the following program: and initial step of 100 V for 5 h, followed by four step gradients of 500 V for 30 min, 500 V for 7 h, 1,000 V to 500 V/h and 8000 V to 13,500 V/h. At the end, a step of 8,000 V to 45,000 V/h was reached. A total of 64,076 V/h were accumulated at the end.

For the second dimension, IPG strips were equilibrated for 15 min in 5 mL of reducing equilibration buffer (75 mM Tris HCl pH 8.8, 6 M urea, 30% (v/v) glycerol, 2% (w/v) SDS, 10 mg/mL of DTT and traces of bromophenol blue) followed by 15 min incubation in 5 mL of alkylating equilibration buffer (75 mM Tris HCl pH 8.8, 6 M urea, 30% (v/v) glycerol, 2% (w/v) SDS, 25 mg/mL of iodoacetamide and traces of bromophenol blue). The second dimension SDS-PAGE was performed on a PROTEAN II xi Basic Electrophoresis System (BioRad) at 1mA/gel and 100V at 15°C, overnight until the bromophenol blue reached the bottom of the gel. The entire electrophoresis unit was protected from direct light during the run.

After electrophoresis, gels were stained for two hours with fluorescent Oriole Solution (BioRad), protected from direct light and under constant shaking.

### Image acquisition and image analysis

Stained gels were immediately observed at excitation/emission wavelengths of 270/604 nm using ChemiDoc MP Imaging System (BioRad). The gel images were captured using ImageLab software, using the automatic exposure and 24.5 cm width.

Images were analyzed with PDQuest Basic Software (Bio-Rad, Hercules, CA, USA). Spots differentially expressed were selected by two different criteria: Qualitative spots (presence/absence), and quantitative spots (*p* = ≤ 0.05) with a fold difference of 2 compared to the control.

### Gel Excision and In-Gel Digestion of proteins

The spots containing the proteins of interest were cut using the EXQuest Spot Cutter (Bio-Rad).

Gel pieces containing the proteins of interest were digested using the DigestPro MS (Intavis) instrument. Proteins were reduced with dithiothreitol (DTT) 10 mM in BCA 50 mM (56°C 1h), alkylated with iodoacetamide (IA) 55 mM in BCA 50mM (30 min) and enzymatically digested with Trypsin Gold Mass Spectrometry Grade (Promega) for 10 h at 37 °C. Peptides were eluted with trifluoroacetic acid (TFA) 0.2% and acetonitrile (AcN) 30%.

### Mass Spectrometry (MS) identification of proteins

Tryptic digests of each spots were desalted with Zip Tip µC18 Pipette Tips (Millipore) according to the recommended protocol and eluted directly on the AnchorChip target plate (Bruker-Daltonics) with α-cyano-4-hydroxycinnamic acid (CHCA) matrix. The MS and tandem MS (MS/MS) spectra were obtained using an UltrafleXtreme MALDI-TOF/TOF mass spectrometer (Bruker-Daltonics) in auto-mode using Flex Control v3.4 (Bruker-Daltonics) and processed using ProteinScape v3.1.3 (Bruker-Daltonics). Peptide spectra were acquired in reflectron mode with 2,000 laser shots per spectrum. Spectra were externally calibrated using peptide calibration standard (Bruker). A total of 2,500 laser shots were accumulated for MS/MS data. The peptide mass fingerprint and MS/MS search were performed on NCBI database for Viridiplantae and *Capsicum annuum* searching for plant proteins. Bacterial proteins of *Microbacterium sp.* 3J1 were searched in databases for *Microbacteriae, Micrococci* and *Actinobacteria.* SwissProt database was used for all organisms, using MASCOT 2.4.0 software integrated together with ProteinScape software (Bruker-Daltonics). The parameters used for the search engine were: monoisotopic peptide mass accuracy of 50 ppm, fragment mass accuracy to ±0.5 Da; maximum of only one missed cleavage; carbamidomethylation of cysteine as a fixed modification and partial oxidation of methionine as a variable modification. There were no restrictions regarding the molecular weight (MW) and isoelectric point (pI) of the proteins. Filtering of peaks was done for known autocatalytic trypsin peaks; the signal to noise threshold was set to 2. The significance threshold was set at *p*< 0.05.

## ACKNOWLEDGMENTS

This work was funded by the Spanish Ministry for Economy and Competitiveness, within the context of the research projects CTM2017-84332-R, and CGL2017-91737-EXP and by the Andalusian Regional Government under the aegis of research project P11-RNM-7844 and PY18-976. Juan I. Vílchez was funded by the Spanish Ministry of Economy with a FPU fellowships.

